# *LTP2* hypomorphs show genotype-by-environment interaction in early seedling traits in *Arabidopsis thaliana*

**DOI:** 10.1101/2023.05.11.540469

**Authors:** Cristina M Alexandre, Kerry L Bubb, Karla M Schultz, Janne Lempe, Josh T Cuperus, Christine Queitsch

## Abstract

Isogenic individuals can display seemingly stochastic phenotypic differences, limiting the accuracy of genotype-to-phenotype predictions. The extent of this phenotypic variation depends in part on genetic background, raising questions about the genes involved in controlling stochastic phenotypic variation. Focusing on early seedling traits in *Arabidopsis thaliana*, we found that hypomorphs of the cuticle-related gene *LTP2* greatly increased variation in seedling phenotypes, including hypocotyl length, gravitropism and cuticle permeability. Many *ltp2* hypocotyls were significantly shorter than wild-type hypocotyls while others resembled the wild type. Differences in epidermal properties and gene expression between *ltp2* seedlings with long and short hypocotyls suggest a loss of cuticle integrity as the primary determinant of the observed phenotypic variation. We identified environmental conditions that reveal or mask the increased variation in *ltp2* hypomorphs, and found that increased expression of its closest paralog *LTP1* is necessary for *ltp2* phenotypes. Our results illustrate how decreased expression of a single gene can generate starkly increased phenotypic variation in isogenic individuals in response to an environmental challenge.

## INTRODUCTION

Genetically identical individuals can develop different phenotypes. Understanding the mechanistic underpinnings of this non-genetic phenotypic variation and the relative contributions of environmental factors and stochasticity holds promise for more accurate genotype-phenotype predictions. As early as 1920, Sewall Wright wrote that a sizable fraction of the environmental contribution to phenotypic variation is likely missed experimentally, and suggested that stochasticity may play an important role in shaping individual phenotypes (Wright, 1920). Individuals sampled from an isogenic population can differ because of (1) stochastic differences in gene expression (Elowitz *et al*., 2002; Blake *et al*., 2003; Raser & O’Shea, 2004; Volfson *et al*., 2006; Lomvardas *et al*., 2006; Gimelbrant *et al*., 2007), protein levels (Feinerman *et al*., 2008) or metabolic states (Smith *et al*., 2007; Heerden *et al*., 2014); (2) parental effects (Perez *et al*., 2017); or (3) differences in microenvironments, relative position or other contextual information (Eagle & Levine, 1967; Snijder *et al*., 2009). None of these possible causes are mutually exclusive. Even subtle differences can have large cumulative effects, because the internal state of individuals affects how they respond to environment factors. A classic example of environmentally-induced heterogeneity is the behavior of a collection of temperature-sensitive cell-cycle mutants in yeast. When grown asynchronously, these mutants show heterogeneous, non-heritable phenotypes due to individual cells experiencing the restrictive temperature treatment at different stages in the cell cycle (Hartwell *et al*., 1974). A multitude of studies in animals and plants have shown that the extent of non-genetic phenotypic variation across individuals and populations depends in part on genotype, with some genetic backgrounds of the same species showing greater non-genetic variation (or less phenotypic robustness) than others (Waddington, 1942; Whitlock & Fowler, 1999; Ros *et al*., 2004; Hall *et al*., 2007; Hill *et al*., 2007; Sangster *et al*., 2008; Ansel *et al*., 2008; Shen *et al*., 2012; Ayroles *et al*., 2015; Katsanos *et al*., 2017).

The more that increased, non-genetic phenotypic variation occurs in a particular genetic background, the lower will be our ability to predict phenotype from genotype, because the same genetic variants will show different expressivity in different individuals (Queitsch *et al*., 2002; Eldar *et al*., 2009; Raj *et al*., 2010; Burga *et al*., 2011; Casanueva *et al*., 2012; Lachowiec *et al*., 2016; Zabinsky *et al*., 2019). Therefore, non-genetic phenotypic variation has wide-ranging implications, from cancer drug resistance (Sharma *et al*., 2010; Shaffer *et al*., 2017; Márquez-Jurado *et al*., 2018; Emert *et al*., 2021) to microbial bioproduction (Delvigne & Goffin, 2014; Xiao *et al*., 2016). In agriculture, trait uniformity is particularly highly prized (Finch-Savage & Bassel, 2016; Tran *et al*., 2017), and breeding programs rely on parental performance. A better understanding of the genetic underpinnings of non-genetic phenotypic variation and their interplay with environmental factors might inform targeted breeding of more robustly performing varieties. In plants, several genes have been identified that affect non-genetic variation of traits like growth (Joseph *et al*., 2015; Illouz-Eliaz *et al*., 2019), organ size or number (Hall *et al*., 2007; Hong *et al*., 2016), germination (Abley *et al*., 2021), early seedling phenotypes (Queitsch *et al*., 2002; Mason *et al*., 2016; Lachowiec *et al*., 2018; Lemus *et al*., 2023) and defense metabolites (Jimenez-Gomez *et al*., 2011; Joseph *et al*., 2015).

Here, we focus on hypocotyl elongation in the dark, an adaptive trait relevant for seedling establishment. Hypocotyl elongation in the dark shows large non-genetic variation in *A. thaliana* (∼10% coefficient of variation in hypocotyl length(Maloof *et al*., 2001; Queitsch *et al*., 2002; Borevitz *et al*., 2002; Sangster *et al*., 2008; Lachowiec *et al*., 2018). We found that hypomorphs of *LTP2* (*LIPID TRANSFER PROTEIN 2*/AT2G38530), a highly expressed gene in dark-grown seedlings, show increased phenotypic variation under specific environmental conditions. Plant Lipid Transfer Proteins (LTPs) are a family of small (∼9 kDa) lipid-binding, cysteine-rich proteins that are commonly found in the shoot epidermis (Kader, 1996; Yeats & Rose, 2008). While structurally similar, LTPs have different expression patterns, suggesting functional specialization (Arondel *et al*., 2000; Chae *et al*., 2010). LTPs have been associated with antimicrobial activity and cuticle physiology, and are implicated in a wide variety of biological processes (Molina & García-Olmedo, 1993; Buhot *et al*., 2001; Maldonado *et al*., 2002; Nieuwland *et al*., 2005; Cameron *et al*., 2006; Debono *et al*., 2009; Chae *et al*., 2009; Potocka *et al*., 2012; Finkina *et al*., 2016; Gao *et al*., 2016).

LTP2 is abundant in the epidermal cell wall of dark-grown hypocotyls (Irshad *et al*., 2008; Jacq *et al*., 2017), where it promotes cuticle integrity and desiccation tolerance (Jacq *et al*., 2017). We found that under certain environmental conditions, *LTP2* is necessary for full hypocotyl elongation in the dark, and that a decrease in *LTP2* expression increases variation in hypocotyl length, gravitropism and cuticle permeability in isogenic seedlings. Differences in epidermal morphology and cuticle permeability between long and short *ltp2* hypocotyls and between growth conditions that promote or mask trait variation suggest that loss of cuticle integrity in *ltp2* hypocotyls is the main determinant of this background’s increased non-genetic phenotypic variation.

## RESULTS

### Decreased *LTP2* expression increases non-genetic phenotypic variation in skotomorphogenesis

Young seedlings grown in the dark show common skotomorphogenic phenotypes with elongated hypocotyls, etiolated cotyledons and short roots (Gendreau *et al*., 1997; Vandenbussche *et al*., 2005). *LTP2* is among the most highly expressed genes in dark-grown shoots (99th percentile, Figure S1A, B), suggesting that its function is required during skotomorphogenesis. To measure the impact of *LTP2* on the phenotypic variation of elongating hypocotyls, we used two *ltp2* hypomorphs, *ltp2-1* and *ltp2-2,* harboring T-DNA insertions less than 500 bp upstream of the *LTP2* transcriptional start site (Figure 1A). Both lines expressed less than 25% of wild-type *LTP2* RNA levels in dark-grown shoots (Figure 1B), and showed similar defects in skotomorphogenesis. When grown in the dark, *ltp2* hypocotyls were shorter (Figure 1C, D) and more variable than wild type (Col-0; Figure 1C), in both length (Figure 1E) and orientation (Figure 1F). Hypocotyl lengths were almost twice as variable in *ltp2-1* as in Col-0 wild type (merged coefficient of variation, CV, of 21% *vs* 13%), with *ltp2-2* being slightly less variable (merged CV 18%; Figure 1E). The higher variation in the hypomorphs was not due to a bimodal distribution but to a continuous, wider distribution of hypocotyl lengths (Figure 1D). An even larger difference was measured for hypocotyl orientation (CV 34-37% *vs* 10%; Figure 1F), a proxy for reduced negative gravitropism. However, there was no substantial correlation between hypocotyl length and orientation of individual seedlings (Figure S1C). Consistent with a non-genetic origin for this increased variation in hypocotyl length, the selfed offspring of *ltp2* parents with either long or short hypocotyls showed similar mean hypocotyl lengths (Figure 1G). Other notable *ltp2* phenotypes included longer roots than hypocotyls, resulting in a smaller hypocotyl/root ratio per individual seedling than in Col-0 wild type (Figure 1H), and a tendency for open and expanded cotyledons (Figure 1C arrowhead, 1I). Taken together, reduced levels of *LTP2* globally affect non-genetic variation in skotomorphogenesis.

**Figure 1.**
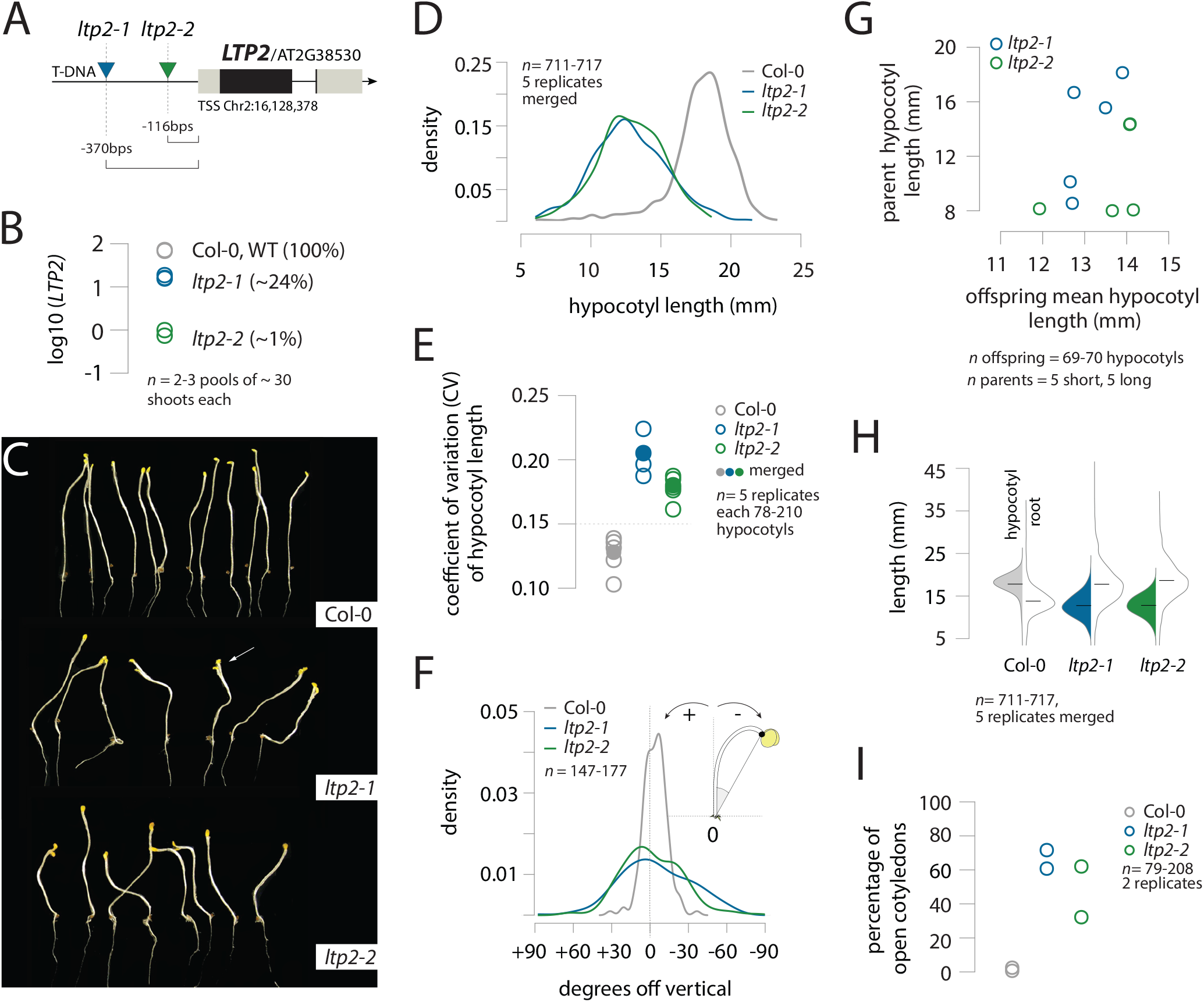
*LTP2* hypomorphs increase phenotypic variation in skotomorphogenesis traits. **A**. Approximate location of the T-DNA insertion (LB) relative to the *LTP2* gene in *ltp2-1* and *ltp2-2* lines. **B**. *LTP2* relative expression in the shoots of wild-type (Col-0), *ltp2-1* and *ltp2-2* seedlings grown in the dark for 7 days (d) after stratification. In parenthesis are the % of Col-0 *LTP2* transcript levels detected in the *ltp2* lines. **C**. Representative image of Col-0 (top), *ltp2-1* (middle) and *ltp2-2* (bottom) dark-grown seedlings at 7 days after stratification; the roots were trimmed. **D**. Density lines of the distributions of hypocotyl length scored at 7 days post-stratification in Col-0 and *ltp2* seedlings grown in MS media with 1% sucrose; each line represents the merged distribution of 5 biological replicates (*n*=78-210 each) per genotype. **E**. Coefficient of variation (sd/mean) in hypocotyl length, a measure of variation, for (1) 5 biological replicates (open circles) and (2) merged for all 5 replicates (filled circle, same data as in **D**). **F**. Hypocotyl negative gravitropism measured as the deviation from vertical (in degrees) of the hypocotyl apex in 7d dark-grown seedlings grown on MS+1% sucrose. The density distribution of measured angles for all genotypes is plotted. **G**. Comparison between the hypocotyl length of individual *ltp2* parents (5 long and 5 short) and the mean hypocotyl length of their offspring; all lengths measured at 7d post-stratification on seedlings grown on MS+1% sucrose. **H**. Beanplots of hypocotyl (left side) and root (right side) length from the merged dataset used in **D** and **E**. **I**. Percentage of dark-grown seedlings with an open cotyledon phenotype, in wild-type (Col-0), *ltp2-1* and *ltp2-2* genotypes, at 7 days after stratification on MS+ 1% sucrose. Two replicates are shown (n=79-208).

### *ltp2* phenotypes show strong gene-by-environment interaction

Next, we explored internal and external factors that might be associated with the hypocotyl length of individual seedlings or modulate the extent of phenotypic variation in *ltp2* hypomorphs. We started by examining the influence of seed batch, seed germination and seed size on hypocotyl length. We found similar mean hypocotyl lengths (Figure S2A) and similarly high coefficients of variation with different *ltp2* seed batches (Figure 1E). There was no germination delay or increased heterogeneity in germination in the *ltp2* hypomorphs relative to Col-0 wild type, as inferred from the correlation of hypocotyl and root length and its time-dependent drop in seedling development (Figure S2B). Finally, hypocotyl length of individual seedlings was not substantially explained by seed size in either the *ltp2* hypomorphs or Col-0 wild type (Figure S2C).

We continued by examining the impact of the growth medium. In the dark, hypocotyls grow longer if provided with sucrose. However, the presence of sucrose also modifies hypocotyl and root growth kinetics (Kircher & Schopfer, 2012) (Figure S3A) and is associated with a decrease in the hypocotyl/root ratio compared to growth conditions without sucrose (Kircher & Schopfer, 2012) (Figure S3B). We reasoned that the presence of sucrose might affect variation in hypocotyl elongation. In our typical experimental setup, seedlings grow in vertical plates, with hypocotyls in direct contact with the medium, and sucrose-driven hypocotyl elongation depends on shoot uptake(Figure S3C). A comparison between *ltp2* seedlings grown on MS media, MS+1% sucrose or MS+1% glucose revealed that *ltp2* phenotypes were strongly sucrose-dependent. Either removing sucrose or replacing it with glucose was sufficient to rescue all *ltp2* growth phenotypes, including differences in mean length and coefficient of variation (Figure 2A,B), hypocotyl negative gravitropism (Figure 2C) and hypocotyl length/root length ratio (Figure 2D). Therefore, *ltp2* hypocotyls can fully elongate under favorable conditions. Sucrose did not cause an increase in the coefficient of variation of hypocotyl length in wild type Col-0, despite a noticeable increase in mean length in this condition (Figure 2A, B).

**Figure 2.**
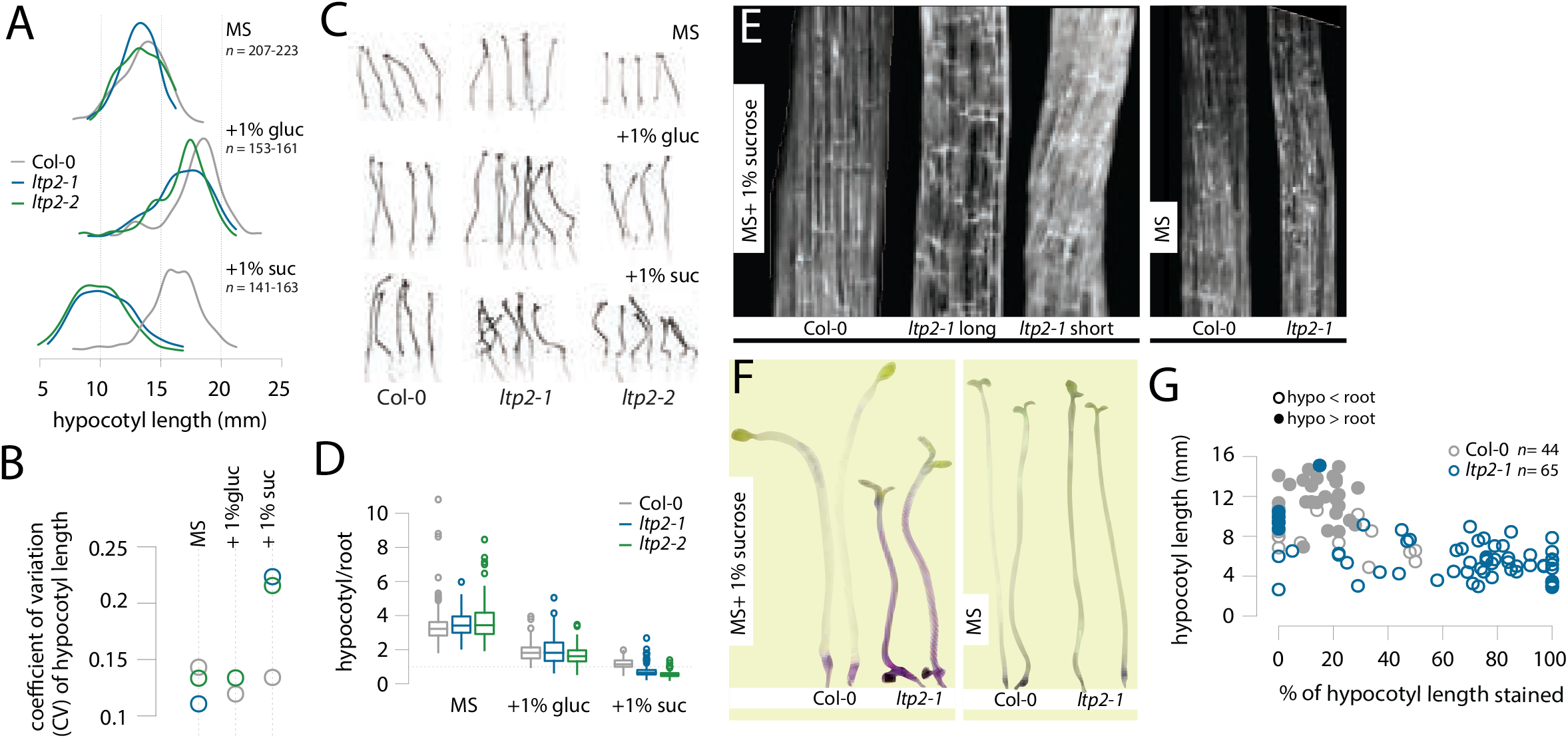
Sucrose-dependent loss of cuticle integrity in *ltp2* is associated with short hypocotyls. **A**. Density lines of the distribution of hypocotyl length for seedlings grown on plates containing MS medium, MS+1% glucose or MS+1% sucrose for 7 days post-stratification. Shown are the merged distributions of two biological replicates per condition/per genotype. **B**. Coefficient of variation in hypocotyl length for the distributions shown in **A**. **C**. Representative shoots of 7d dark-grown seedlings grown on plates containing MS medium, MS+1% glucose or MS+1% sucrose. Hypocotyl negative gravitropism is largely rescued in *ltp2* seedlings grown without sucrose. **D**. Boxplots comparing the hypocotyl/root ratio for every seedling in **A** across different growth media. **E**. Magnified images of the epidermal surface of Col-0 and *ltp2-1* hypocotyls at 7d post-stratification when grown with (left) or without (right) 1% sucrose. **F**. Comparing the effects of 1% sucrose on hypocotyl cuticle permeability: dark-grown seedlings of Col-0 and *ltp2-1* were stained with toluidine blue after growing for 5 days on MS+1% sucrose (left) or 7 days on MS alone (right). **G**. Relationship between hypocotyl length at 7 days after stratification on MS+1% sucrose and toluidine blue staining coverage expressed in % of hypocotyl length. Filled dots represent seedlings with the wild-type developmental pattern of hypocotyl length > root length.

We wondered whether the effects of sucrose on *ltp2* seedlings involved osmotic stress. To test this possibility, we added mannitol to the growth medium. Mannitol causes strong osmotic stress in *Arabidopsis* seedlings (Zwiewka *et al*., 2015; Kalve *et al*., 2020). Adding equimolar amounts of mannitol (29mM) and sucrose (1%) together improved, rather than aggravated, the *ltp2* phenotypic defects (Figure S4). We also considered whether *ltp2* seedlings were deficient in sucrose uptake but concluded that this scenario is unlikely because of the following results: The hypocotyls of *ltp2* seedlings remain sucrose-sensitive and show similar sucrose-driven responses as those of wild-type (Figure S5). Further, doubling the amount of sucrose inhibited hypocotyl elongation to the same extent in *ltp2* and wild-type seedlings (Figure S5D). Lastly, *ltp2* hypocotyls were shorter when grown on MS+1% sucrose compared to MS alone (Figure 2A), with a significant difference in mean values (*ltp2-1*: 3.03mm CI95% 2.62-345, *ltp2-2*: 3.96mm CI95% 3.54-4.38, *ltp2-1*: t(258.75)=14.459; p <2.2e-16, *ltp2-2*: t(274.64)=18.5; p <2.2e-16), inconsistent with deficient sucrose uptake. We conclude that neither osmotic stress nor sucrose uptake contribute substantially to the *ltp2* phenotypes.

### A sucrose-dependent increase in cuticle permeability is associated with short *ltp2* hypocotyls

A comparison of *ltp2* hypocotyls grown with and without sucrose revealed epidermal features that were associated with short hypocotyls. Compared to wild-type Col-0, the epidermal surface of *ltp2* hypocotyls was not smooth, and instead was fuzzy or wrinkled (Figure 2E); this *ltp2* phenotype was sucrose-specific and much stronger in short than in long *ltp2* hypocotyls (Figure 2E). Furthermore, growth with sucrose greatly increased permeability to the water-soluble dye toluidine blue, an indicator of cuticle integrity (Tanaka *et al*., 2004), in *ltp2* hypocotyls, but not in Col-0 hypocotyls, compared to controls without sucrose. Most *ltp2* hypocotyls stained deeply, and over more than 50% of their full length (Figure 2F, G), consistent with reduced cuticle integrity (Jacq *et al*., 2017). The extent of staining varied widely in *ltp2* hypocotyls, from 0 to 100% (Figure 2G). On average, heavily stained hypocotyls were smaller than those not stained (Figure 2G), and *ltp2* seedlings with the wild-type-like hypocotyl length/root length ratio > 1 were less stained overall (Figure 2G). In contrast, staining in Col-0 wild-type hypocotyls was generally weak and spatially restricted (Figure 2F, G). These results suggest that altered cuticle integrity contributes to the sucrose-dependent inhibition of hypocotyl elongation in *ltp2* seedlings.

### Hypocotyl length is associated with many transcriptional differences in *ltp2* seedlings

To identify what molecular functions are associated with non-genetic variation in hypocotyl length, we compared the transcriptional profiles of Col-0 and *ltp2-1* seedlings with short (S) and long (L) hypocotyls (bottom and top 15th percentiles, respectively; Figure 3A). For each genotype by length combination, we performed bulk RNA-seq on two replicate pools of 70 shoots each (Figure 3A; see Figure S6A-C for QC metrics). Principal component analysis showed three clearly distinguishable clusters: one formed by Col-0 L and S samples, a second formed only by *ltp2-1* L samples and a third containing the *ltp2-1* S samples (Figure 3B). Nearly half of the global variance in gene expression (47%, PC1) was correlated with mean hypocotyl length (Figure 3C). Over 20 times as many differentially expressed genes (DEGs, log2FC >=1 and p-adj < 0.01) were found between the short and long *ltp2* samples (584, Table S2) compared to the Col-0 samples (24, Table S1). The differentially expressed genes associated with hypocotyl length in *ltp2* samples were enriched in gene ontology (GO) terms related to cell wall modification, response to stress and plant defense (Figure 3D, fold-enrichment >= 2, p-adj < 0.05). *ltp2* samples with short hypocotyls showed downregulation of cell wall-related genes that promote growth, like *PGX1* and *XTH20* (Miedes *et al*., 2013; Xiao *et al*., 2014), secondary cell wall biosynthesis genes, including the three main laccases *LAC4*, *LAC11* and *LAC17,* and several peroxidases and genes related to casparian strip deposition (Figure 3E). Although casparian strip biology is not well studied outside of roots, the casparian strip is present in dark-grown hypocotyls (Karahara, 2012; Geldner, 2013). None of the genes in the enriched term “casparian strip” were differentially expressed between Col-0 wild-type and *ltp2* seedlings with long hypocotyls (Figure 3E), suggesting that the down-regulation of these genes may contribute to the phenotype of *ltp2* seedlings with short hypocotyls.

**Figure 3.**
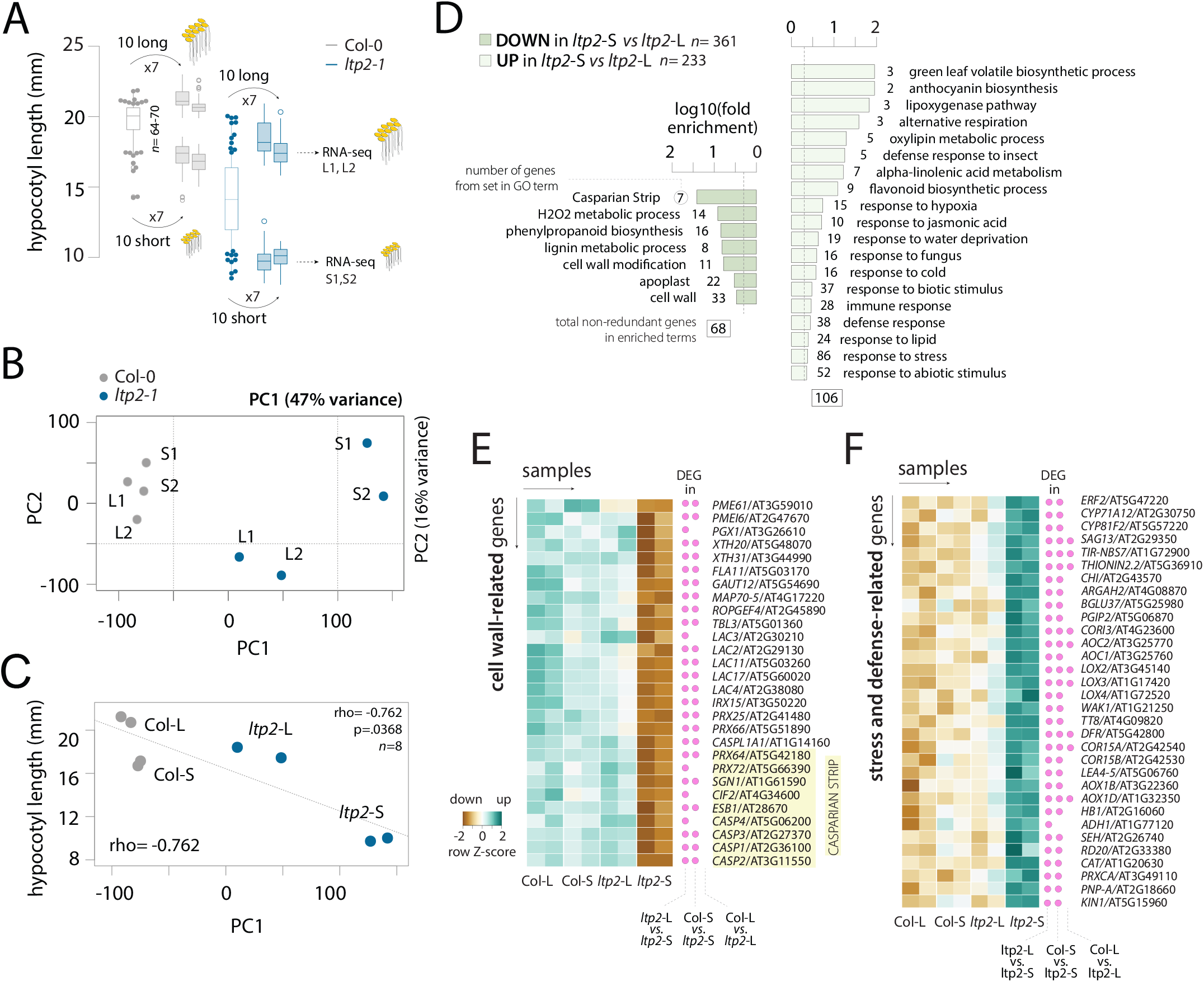
Hypocotyl length is associated with many transcriptional differences in *ltp2* but not Col-0 seedlings, mostly related to stress, defense and growth. **A.** Experimental setup for collecting RNA-seq samples. For Col-0 and *ltp2-1*, the first boxplot shows the distribution of hypocotyl length from one out of seven hypocotyl assays done per replicate; the second set of four boxplots (two replicates with short and two replicates with long hypocotyls) shows the actual distribution of hypocotyl lengths (n=70) from the selected individuals used to generate RNA-seq libraries. **B**. Biplot of Principal Component Analysis showing the first two PCs. The percentage of the total variance explained by PC1 and PC2 is indicated on the top right corner. **C**. Correlation between mean hypocotyl length for the eight RNA-seq samples and PC1 loadings; shown on the bottom left corner is the Spearman rho. **D**. Shortlisted GO enrichments (fold-enrichment >=2 and adjusted p-value < 0.05) for the set of 584 differentially expressed genes between *ltp2*-L and *ltp2*-S, split by down- and upregulated genes. Shown are the log10(fold-enrichment) and the number of DEGs per enriched term. **E-F** Row-scaled heatmap visualizations of a subset of DEGs downregulated (**E**) or upregulated (**F**) in *ltp2*-S relative to *ltp2*-L. Pink dots indicate whether each gene was also a DEG on other pairwise-comparisons.

We also observed the upregulation of a variety of stress and defense-response genes related to hypoxia, response to fungus, anthocyanin and jasmonate biosynthesis (Figure 3F). A few of these genes (11), including several of the most highly upregulated genes in *ltp2* seedlings with short hypocotyls (Figure S6E), were also upregulated in Col-0 seedlings with short hypocotyls compared to Col-0 seedlings with long hypocotyls (Figure S6D). This result supports the idea that the phenotypic impact of perceived stress on hypocotyl length for a given individual can be genotype-independent; however, *ltp2* individuals experience this impact far more frequently and far more severely.

The majority of the differentially expressed genes (474, 81%) between the long and short *ltp2* samples were also differentially expressed between the short samples of *ltp2* and Col-0. This result likely reflects that the difference in mean hypocotyl length between the short *ltp2* and the short wild-type samples is about as large as the difference between the long and the short *ltp2* samples (Figure S6F). However, we found more than twice as many differentially expressed genes (1638) in the comparison of the short *ltp2* and short wild-type samples (Figure S6G).

These differentially expressed genes showed similar gene ontology enrichments as found for the comparison of long and short *ltp2* samples, related to hypoxia, oxidative stress, plant defense and cell wall metabolism, with additional growth-related terms such as response to auxin (Figure S6H). Indeed, we found that many auxin-responsive genes, including some with known roles in hypocotyl elongation and/or gravitropism like *SAUR19/23/24/32* (Park *et al*., 2007; Spartz *et al*., 2012), *ARGOS* (Rai *et al*., 2015), *SHY2* (Reed *et al*., 1998; Tian *et al*., 2002) and *HAT2* (Sawa *et al*., 2002), were downregulated in *ltp2* seedlings with short hypocotyls compared to wild-type seedlings with short hypocotyls, but not compared to *ltp2* seedlings with long hypocotyls (Table S5). Most of the differentially expressed genes in the short sample comparison (996/1164, 86%) did not overlap with the differentially expressed genes in the long sample comparison. Because the *ltp2* seedlings with long hypocotyls and Col-0 wild-type seedlings with long hypocotyls differ in genotype but little in mean hypocotyl length, we conclude that the gene expression differences across samples are strongly associated with hypocotyl length, in particular with the strongly reduced hypocotyl length of short *ltp2* seedlings.

### *ltp2* phenotypes depend on upregulation of its closest paralog *LTP1*

Focusing on cuticle-related genes (see Methods; from (Li-Beisson *et al*., 2013)), we found that 21 out of the 160 expressed in dark-grown shoots were deregulated in *ltp2* relative to Col-0 seedlings (30% more than expected, p=0.019; Table S6). Most of these genes (19/21) were upregulated in *ltp2*, including six other LTP paralogs (Figure 4A). To explore to what extent close paralogs may compensate for reduced *LTP2* expression, we examined the role of its nearest paralog, *LTP1* (Figure 4B, Figure S7A). Analysis of *LTP1* and *LTP2* expression in light- and dark-grown wild-type seedlings showed a reciprocal expression pattern that suggests non-redundant roles during hypocotyl elongation: whereas *LTP1* was about 100 times more abundant than *LTP2* in the light, it was 1/10 times as abundant in the dark (Figure 4C). In dark-grown seedlings, *LTP2* accounts for more than 86% of global Type I LTP expression compared to only 5% for *LTP1* (Figure S7B). In stark contrast, *LTP1* tends to be the more abundantly expressed paralog in many other conditions, organs, and developmental stages (41/56 samples; Figure S7C). Moreover, we found that *LTP1* expression was no longer repressed in the dark in *ltp2* seedlings (Figure 4C, Figure S7B). To rule out that this trend was an artifact of bulk expression analysis, we measured *LTP1* and *LTP2* expression in individual wild-type Col-0 and *ltp2-1* seedlings and measured the length of their hypocotyls. *LTP1* expression levels in individual *ltp2-1* seedlings recapitulated our bulk observations, and their hypocotyl lengths were negatively correlated with *LTP1* expression (Figure S8; Spearman rho = -0.401; p=.0108). We further found that *LTP1* and *LTP2* expression levels were correlated across individual seedlings in both wild-type and *ltp2-1* seedlings (Figure S8A; Col-0: 0.832; p< 2.2e-16; *ltp2-1*: 0.933; p<2.2e-16), suggesting that the *LTP1/LTP2* expression ratio may matter for phenotype. Indeed, we observed that *LTP1/LTP2* expression ratios were weakly but significantly correlated with hypocotyl length across individual *ltp2-1* and wild-type seedlings (Figure S8E; Spearman’s rho = 0.484, p = 0.0018 *ltp2-1*; Spearman’s rho = 0.437, p= 0.0059 Col-0).

**Figure 4.**
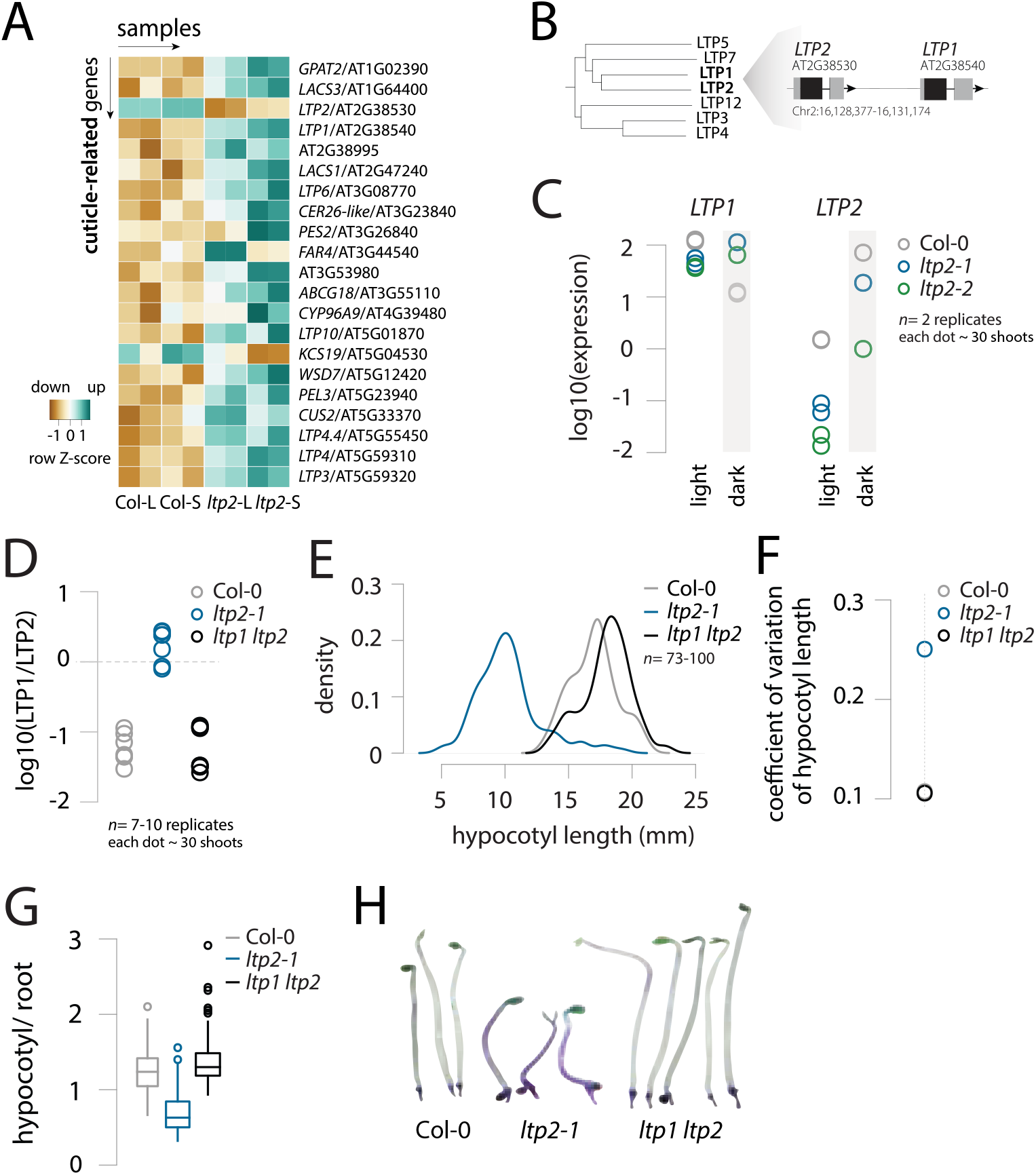
The *ltp2* phenotype depends on the expression ratio of *LTP2* and its close paralog *LTP1*. **A**. Row-scaled heatmap visualization of cuticle-related genes differentially expressed between Col-0 and *ltp2* samples. **B**. Detail of a tree depicting amino acid sequence similarity among PR14/Type I LTP proteins (full tree in Figure S5A). **C**. Relative expression of *LTP1* and *LTP2* in the shoots of light or dark-grown (gray box) Col-0 and *ltp2* seedlings grown for 7 days after stratification on MS+ 1% sucrose. **D**. Comparison of the *LTP1/LTP2* expression ratio between Col-0, *ltp2-1* and an *ltp1ltp2* double mutant. **E-G** Hypocotyl length distribution (**E**), coefficient of variation of hypocotyl length (**F**) and (**G**) hypocotyl/root ratio for Col-0, *ltp2-1* and an *ltp1 ltp2* double knockdown; all seedlings were grown in the dark for 7d after stratification on MS+ 1% sucrose. **H**. Representative images of toluidine blue stained etiolated hypocotyls from Col-0, *ltp2-1* and an *ltp1 ltp2* double mutant grown for 7 days after stratification on MS+ 1% sucrose

We next tested whether the altered expression ratio of the paralogs might contribute to the *ltp2* phenotypes in transgenic lines. We used an artificial microRNA to simultaneously knock-down *LTP1* and *LTP2* gene expression while aiming for a similar *LTP1*/*LTP2* ratio as found in Col-0 wild-type (Figure 4D). Indeed, the mean value and variation of hypocotyl length of the *ltp1 ltp2* double knock-down seedlings closely resembled those of Col-0 wild-type seedlings with both distributions largely overlapping (Figure 4E, F). The *ltp1 ltp2* double knock-down seedlings were also similar to Col-0 wild-type seedlings in hypocotyl/root ratio (Figure 4G) and hypocotyl cuticle integrity (Figure 4H). This result is even more remarkable considering that the expression levels of the individual *LTP1* and *LTP2* genes differed substantially in the *ltp1 ltp2* double knock-down seedlings from those observed in wild-type Col-0 seedlings (Figure S9), consistent with their expression ratio as a determinant of the hypomorph phenotype. We conclude that the upregulation of *LTP1* compared to low expression levels of *LTP2* in dark-grown seedlings contributes to the variable skotomorphogenesis phenotypes of the *ltp2* hypomorphs.

## DISCUSSION

Here, we show that the gene *LTP2* plays an important role in shaping skotomorphogenesis traits. Hypomorphs of *LTP2* show increased phenotypic variation in three hypocotyl traits: negative gravitropism, length and cuticle permeability. For the latter two, the increased variation was accompanied by altered mean values relative to wild type, with most *ltp2* dark-grown seedlings having short hypocotyls with highly permeable cuticles. Variation in hypocotyl length increases in dark-grown hypocotyls upon perturbation of the chaperone Hsp90 (Queitsch *et al*., 2002; Sangster *et al*., 2008), or its client protein BEH4 (Lachowiec *et al*., 2018), as well as in *AGO1* hypomorphs (Lemus *et al*., 2023). As for the *ltp2* hypomorphs described here, in these three cases, the increased variation in hypocotyl length was also accompanied by decreased length means. This concordance of changes in mean and variation might be expected if wild-type hypocotyls reach lengths close to their maximum physiological limit under these experimental conditions.

In *ltp2* hypocotyls, a strong defect in cuticle permeability was associated with reduced elongation and increased phenotypic variation compared to wild type; however, these phenotypes depended on exposure to sucrose. This remarkably strong gene-by-environment effect is unlikely due to a higher requirement for LTP2 function, as hypocotyls elongate more when provided with an exogenous carbon source. *LTP2* is required in dark-grown hypocotyls to seal the cuticle and prevent water loss (Jacq *et al*., 2017). Moreover, increased cuticle permeability is linked to structural defects and poor adhesion between the cuticle and the rest of the cell wall. Our finding that *ltp2* hypocotyls were much more susceptible to sucrose-induced dye uptake than wild type is consistent with *LTP2*’s role in cuticle sealing and suggests that sucrose acts as a cuticle stress. Sucrose may trigger gene expression changes that modify cuticle composition or act directly to increase cuticle hydration, the mechanical strain associated with water accumulating in the cuticle and in gaps between the cuticle and the cell wall. This added strain could lead to further cuticle detachment in sensitized *ltp2* hypocotyls, thus aggravating their documented cuticle integrity defect (Jacq *et al*., 2017). The sucrose-induced loss of cuticle integrity is likely the primary determinant of the shorter *ltp2* hypocotyls and their increased variation in length.

Dark-grown long hypocotyls have thick cuticles (Gendreau *et al*., 1997), and shorter hypocotyls are associated with pharmacological or genetic disruptions in cuticular wax biosynthesis and deposition (Narukawa *et al*., 2016). The greater need for cuticle integrity during hypocotyl elongation in the dark may reflect a seedling’s push upwards through the soil while minimizing abrasion and the need for structural reinforcement due to cell wall thinning in very long cells (Derbyshire *et al*., 2007). Loss of cuticle barrier function causes water loss and activates cuticle-dependent defense priming (Chassot *et al*., 2007; Bessire *et al*., 2007; L’Haridon *et al*., 2011; Serrano *et al*., 2014), which inhibits growth, and is consistent with the gene ontology enrichments observed among upregulated genes in *ltp2* seedlings with short hypocotyls compared to those with long ones. However, the precise mechanisms by which *LPT2* facilitates the elongation of dark-grown hypocotyls remain unknown.

In contrast to phenotypic buffers, which act on gene regulation (Lachowiec *et al*., 2016, 2018; Lemus *et al*., 2023) or protein folding (Queitsch *et al*., 2002; Sangster *et al*., 2008; Lachowiec *et al*., 2016; Zabinsky *et al*., 2019), *LTP2* appears to act structurally by sealing the cuticle. The integrity of this seal determines the effectiveness of the cuticle as an insulating barrier, thus providing a simple mechanism of phenotypic robustness against environmental insults.

Biophysical and regulatory complexity makes plant cuticles liable to harbor considerable non-genetic variation in their composition, ultra-structure and properties, in particular when considering the complexity of the environments plants face. Upon damage or genetic perturbation, this intrinsic variation is amplified as shown by the disorganized cuticles of mutants with cutin defects or increased cuticle permeability (Lolle *et al*., 1992; Sieber *et al*., 2000; Wellesen *et al*., 2001; Chen *et al*., 2003; Schnurr *et al*., 2004; Kurdyukov *et al*., 2006; Takahashi *et al*., 2010).

Loss of cuticle integrity may not fully explain the increased phenotypic variation of *ltp2* seedlings. We cannot rule out that the increased phenotypic variation depends on another, yet undiscovered *LTP2*- function. However, our results that show genetic interaction between the close paralogs *LTP1* and *LTP2*, and that their expression ratio is a determinant of increased cuticle permeability and phenotypic variation during skotomorphogenesis, strongly support our interpretation. In crown gall tumors, the only other context outside of skotomorphogenesis where *ltp2* phenotypes have been identified (Jacq *et al*., 2017), *LTP1* is also highly upregulated (Deeken *et al*., 2016). This finding is consistent with our result that the upregulation of *LTP1* compared to *LTP2* expression is predictive of the deleterious *ltp2* phenotypes.

While paralogs with redundant or partially redundant functions can confer genetic robustness (Gu *et al*., 2003; Kafri *et al*., 2005; DeLuna *et al*., 2008, 2010; Dean *et al*., 2008; Diss *et al*., 2014), this is not universally observed (Ihmels *et al*., 2007; DeLuna *et al*., 2010; Diss *et al*., 2017; Dandage & Landry, 2019). In fact, incomplete functional compensation can be a source of increased phenotypic variation (Burga *et al*., 2011; Bauer *et al*., 2015). For example, *BEH4*, the earliest diverged member of the *BZR/BEH* family of transcription factors, governs phenotypic robustness of hypocotyl length by integrating regulatory cross talk among the six gene family members (Lachowiec *et al*., 2018). Thus, even among these closely related, partially redundant paralogs, increased trait variation arises when the activity of *BEH4* is lost. The loss of properly integrated regulatory cross talk as a cause of increased phenotypic variation is consistent with our findings that upregulation of *LTP1* in the dark is associated with the *ltp2* phenotypes.

Subtle changes in gene expression that percolate through gene regulatory networks and amplify each other to affect expression of certain core genes are thought to underlie complex diseases and complex traits in humans (the omnigenic model, (Boyle *et al*., 2017; Liu *et al*., 2019)). The genetic variants found to be associated with complex diseases and traits in genome-wide association studies (GWAS) typically reside in regulatory regions, likely resulting in hypomorphs. The trait heritability explained by GWAS variants tends to be small, and these variants have little power to predict the disease risk of individuals (Manolio *et al*., 2009; Eichler *et al*., 2010; Gibson, 2012; Khera *et al*., 2018). The low power to predict phenotype from genotype is consistent with high non-genetic trait variation (Queitsch *et al*., 2012). We speculate that this non-genetic variation arises because regulatory variants cause small expression changes that are integrated differently among individuals. In turn, these differences in integrating expression changes will sensitize certain individuals but not others to environmental factors, resulting in different phenotypes. At least for *LTP2*, this interpretation holds: the upregulation of *LTP1* is not sufficient for the observed phenotypes as all *ltp2* seedlings exhibit it. Likewise, all *ltp2* seedlings experience exposure to sucrose; however, not all seedlings have short hypocotyls and show loss of cuticle integrity. The loss of barrier (*i.e*. cuticle) function in this plant example likely holds lessons for studies of human traits and diseases and points to genotype-by-environment effects as a major contributor to non-genetic variation.

Our study highlights that even highly inbred, *de facto* homozygous genetic backgrounds maintain a physiologically relevant reservoir of phenotypic variation, which can be exposed by stress. While stress often increases phenotypic variation in isogenic and inbred populations (Thattai & van Oudenaarden, 2004; Newman *et al*., 2006; Braendle & Félix, 2008; Tokatlidis *et al*., 2010; Uyttewaal *et al*., 2012; Holland *et al*., 2014; Mitosch *et al*., 2017; Sandner *et al*., 2021; de Groot *et al*., 2022), phenotypic robustness (*i.e*., low non-genetic variation) is associated with stress tolerance and vigor in crops and lifestock (Tollenaar & Lee, 2002; Blasco *et al*., 2017; Elgersma *et al*., 2018). A common strategy to achieve phenotypic robustness coupled with high performance in agricultural settings has been the use of F1 hybrids, which are often more uniform in phenotype than their inbred parental lines (Lewis, 1953; Smith *et al*., 1955; Phelan & Austad, 1994). A better understanding of the mechanistic underpinnings of non-genetic phenotypic variation might lead to the development of crops and livestock that combine uniformity of phenotype with broad stress tolerance. This better understanding of non-genetic phenotypic variation will also facilitate efforts to unravel the complexity of non-Mendelian human traits and diseases.

## MATERIALS AND METHODS

### Plant Material

*Arabidopsis thaliana*, all lines are in the Col-0 background. *ltp2-1* is SALK_026257 (ABRC), previously described (Jacq *et al*., 2017). *ltp2-1* plants homozygous for the T-DNA insertion (Chr2:16,128,007 SALK project) were confirmed by PCR analysis with primer pairs CA340 (Chr2:16,128,062) + CA103 (Chr2:16,128,340) and CA249 (Chr2:16,127,7702) + CA103 (primer sequences in Table S7). *ltp2-2* is line DT7-3 (Marjorie Matzke lab) previously described (Kanno *et al*., 2004); contains a transgene insertion located upstream of *LTP2*, mapped in Kanno et al., 2004. Approximate location is Chr2:16,128,261, determined by sequencing the PCR fragment CA340+CA103. *ltp2-2* plants were confirmed homozygous for the transgene insertion by PCR genotyping with the same primers as *ltp2-1*. The *ltp1ltp2* double knock-down line was generated by Agrobacterium-mediated (GV3101) co-transformation of Col-0 plants with the helper plasmid pSOUP (CD3-1124, ABRC) and the plasmid CSHL_0103F2 (ABRC), containing an artificial microRNA against both *LTP2* and *LTP1.* T0 seeds were selected on 15mg/ml phosphinotricin (PPT/BASTA); PPT-resistant T1 plants were then propagated on 25mg/ml PPT; only generations T3 and above, homozygous for the transgene, were used in hypocotyl assays.

### Hypocotyl Assays

1. *Hypocotyl and root length:* Unless otherwise stated, all seed batches were single-seed descent. Seeds were sterilized, resuspended in 0.1% (w/v) Bacto agar (BD, Diagnostics) and spotted in a well-spaced fashion onto square plates containing 1x MS media (Murashige & Skoog Basal Salt Mixture) pH 5.8, with 0.5g/l MES (Sigma-Aldrich), 0.3% Phytagel (Sigma-Aldrich) and 1% sucrose (w/v). Plates were double-sealed with Micropore surgical tape (3M) and stratified in the dark for 4 days at 4C. After stratification, plates were exposed to 3 hours of light, then placed in vertical racks, wrapped in aluminum foil and kept at 22C in a Conviron chamber (50% RH; 16h light/8h dark; ∼ 100 μmol m2 s) for 7 days. For certain experiments, the age and media composition varied as indicated elsewhere. Genotype placement was randomized across racks, and each genotype was distributed over multiple plates per experiment. At 7d after stratification, plates were opened and imaged on a fixed stand. ImageJ was used to trace and measure hypocotyl and root length for every seedling. The hypocotyl was scored from the collet (hypocotyl/root transition zone) till the shoot apical meristem. Root length was used to infer late germination; outlier hypocotyls with < 5mm (at 7 days) were removed from the analysis. Despite our best efforts, hypocotyl elongation proved highly sensitive to random environmental effects, and differences of a few millimeters in mean were not uncommon between replicates, even when controlling for seed batch (see Figure S2A)
2. *Hypocotyl negative gravitropism*: The angle between the shoot apical meristem and an imaginary vertical line, drawn starting at the base of the hypocotyl, was used to quantify the negative gravitropic response in dark-grown hypocotyls at 7d post-stratification. Data was expressed as deviations off vertical (in degrees), with vertical representing perfect negative gravitropism. Measurements were done on ImageJ, as described for hypocotyl length.
3. *Hypocotyl* e*pidermal imaging*: The epidermal surface of dark-grown hypocotyls was imaged at 7d post-stratification. Whole seedlings were placed directly on a glass slide and the hypocotyl imaged using a Zeiss Axioplan microscope at 50x magnification.
4. *Toluidine Blue Assay* Protocol based on Tanaka et al., 2003 but with a lower concentration of dye to avoid saturation in *ltp2*. Whole dark-grown seedlings were immersed in an aqueous solution of 0.02% (w/v) toluidine blue (Sigma) for 2mins with gentle shaking, then washed 3x with distilled water and left in water until imaging. ImageJ was used to determine hypocotyl length and the fraction of the total length stained with toluidine blue.

### Seed Size

Matched datasets of seed size and hypocotyl length were obtained by imaging the same plate twice: (1) after stratification, when seeds were fully imbibed, and (2) after 7days of growth in the dark. The position of the seed and the hypocotyl are the same in both pictures. Between 10-20 seeds were spotted per plate and several plates were employed per genotype. For imaging seeds, pictures were taken under a stereomicroscope, at 5x magnification. Seed area was measured in ImageJ using > 8-bit > Threshold (auto) > Analyze Particles, with settings: area, 5000-infinity; circularity >= 0.7.

### qRT-PCR

Seedling samples were either single individuals or pools of about 30 individuals. Unless otherwise stated, only the shoot was harvested (includes hypocotyl, cotyledons and shoot apical meristem). For roots, rosette leaves and flowers, all samples were pools; roots were excised from 7d old seedlings; rosette leaves and flowers from mature plants. RNA was extracted from LN2 frozen material using TRIZOL (Invitrogen) and DNase I treated. cDNA was synthesized from 250-500mg of total RNA with the RevertAid First Strand cDNA Synthesis kit (Thermo Scientific). qRT-PCR was performed with LightCycler 480 SYBR Green I Master Mix (Roche). Relative gene expression was calculated as 2^-(Ct target - Ct reference), with either *AP2M*/AT5G46630, *UBC21*/AT5G25760 or *PP2A*/AT1G13320 as reference genes. Primers are listed in Table S7.

### RNA-seq data

1. *Setup and sequencing*: For each replicate of Col-0 and *ltp2-1*, seven hypocotyl assays were set up in parallel, each containing 64-70 seedlings. At day 7 after stratification, hypocotyl length was scored for each of the seven assays and from each, only the shoots of the 10 most extreme seedlings at either tail of the distribution (bottom and top 15^th^ percentile) were collected as “short” and “long” samples, respectively. Each sample contained a total of 70 shoots, and corresponds to one replicate. The process was repeated to obtain a second biological replicate, in a total of eight samples (Col-L1, Col-L2, Col-S1, Col-S2, *ltp2*-L1, *ltp2*-L2, *ltp2*-S1, *ltp2*-S2). RNA was extracted from all samples in parallel using the SV Promega Total RNA System followed by NaCl/EtOH precipitation. Generation of RNA-seq libraries, multiplexing and sequencing was outsourced to GENEWIZ (GENEWIZ Inc., NJ). Each library was sequenced on four lanes and two flowcells of an Illumina HiSeq2500 in a 1x50bp SE format.
2. *Pseudoalignment of reads and estimation of transcript abundance* was done using Kallisto (Bray *et al*., 2016).. The kallisto index was built with Arabidopsis thaliana TAIR10 cDNA models ftp://ftp.ensemblgenomes.org/pub/plants/release-50/fasta/arabidopsis_thaliana/cdna/Arabidopsis_thaliana.TAIR10.cdna.all.fa.gz. Transcript abundances were quantified with parameters –single -l 200 -s 20. Kallisto abundance files were parsed into R (v. 3.6.1) using tximport(). The tx2gene file and the TxDb object used TAIR10 annotations ftp://ftp.arabidopsis.org/home/tair/ Genes/TAIR10_genome_release/TAIR10_gff3/TAIR10_GFF3_genes_transposons.gff. The output was a matrix of estimated counts with 26923 rows (genes) and 8 columns (samples).
3. *Differentially Expressed Genes (DEGs)* After non-specific filtering to remove all non-expressed genes (zero counts across all samples; 2,917 genes), and all genes which did not have at least 1 count in all samples (4,023), a filtered matrix of 19,983 genes x 8 samples was converted to integers and used as input for DESeq2 (Love *et al*., 2014), with parameters: condition = genotype x hypocotyl length, with 2 biological replicates per condition. Only genes with a log2FC >=1 & adjusted p-value <= 0.01 were called as DEGs. As high fold-changes are more frequent in genes with low baseline expression, we specifically chose a less stringent fold-change cut-off to avoid discarding genes with very high expression like *LTP2.* DEGs were obtained for four different pairwise comparisons: (1) Col-L *vs* Col-S, (2) Col-L *vs ltp2*-L, (3) Col-S *vs* ltp2-S and (4) *ltp2*-L *vs ltp2*-S (Tables S1-S4).
4. *Principal Component Analysis (PCA)* prcomp() was applied to a transposed standardized matrix of log10(counts) with 19,983 rows x 8 columns.
5. *Gene Ontology (GO) enrichments* were obtained with gProfiler (Raudvere *et al*., 2019) https://biit.cs.ut.ee/gprofiler/gost, with the input to DESeq2 (a list of 19,983 genes) as background. Significant GO terms were shortlisted if fold-enrichment >= 2 & Bonferroni corrected p-value < 0.05. Query genesets were (1) the list of 584 DEGs between *ltp2*-L and *ltp2*-S and (2) the list of 1164 DEGs unique to the comparison Col-S and *ltp2*-S.
6. *Cuticle genes* From the dataset in Li-Beisson et al.,(2013) we selected only genes with annotated roles in cutin or wax biosynthesis and/or deposition (groups Cuticle Synthesis & Transport 1, Fatty Acid Elongation & Wax Biosynthesis and Fatty Acid Synthesis). The curated list contained 224 genes (Table S6), of which 160 were present in our RNA-seq dataset (Table S6), and 21 were differentially expressed between Col-0 and *ltp2* (union set between Col-L vs *ltp2*-L and Col-S vs *ltp2*-S, 1717 DEGs); the hypergeometric test indicates this is a modest over-representation (p= 0.019).

### Protein sequence tree of PR-14/LTP Type I proteins

Protein sequences for PR-14/Type I LTP proteins (Arondel12 *et al*., 2000) were retrieved from NCBI: NP_181388.1 LTP1, NP_181387.1 LTP2, NP_568905.1 LTP3, NP_568904.1 LTP4, NP_190728.1 LTP5, NP_187489.1 LTP6, NP_973466.1 LTP7, NP_179428.1 LTP8, NP_179135.2 LTP9, NP_195807.1 LTP10, NP_680758.3 LTP11, NP_190727.1 LTP12, NP_001078707.1 LTP13, NP_001078780.1 LTP14, NP_192593.3 LTP15. Multiple sequence alignment was done with ClustalW in the msa package (Bodenhofer *et al*., 2015) and the neighbor-joining tree with the ape package (Paradis *et al*., 2004).

### *LTP1* and *LTP2* global expression pattern

We used the Digital Expression Explorer 2 repository (Ziemann *et al*., 2019) https://dee2.io/ to retrieve uniformly processed RNA-seq data from Arabidopsis thaliana. We curated a dataset of 56 samples, spanning several organs, contexts and developmental stages (Table S8), and kept only genes with at least 3 counts in one sample out of the 56 (26,372 genes x 56 samples).

## DATA ACCESS

RNA-seq data is available at GEO (https://www.ncbi.nlm.nih.gov/geo/) BioProject ID PRJNA856271.

## ACKNOWLEDGEMENTS

This work was supported by the National Science Foundation (NSF MCB-1242744, RESEARCH-PGR grant 1748843, PlantSynBio 2240888 to C.Q.) and the National Institute of Health (NIH NHGRI 5RM1HG010461, NIH NIGMS R35GM139532 to C.Q.)

## FIGURE LEGENDS

**Figure S1. A**. *LTP2* is induced in the dark. Shown is the relative expression of *LTP2* in light vs dark-grown shoots at 7d post-stratification (MS+1% sucrose). B. Boxplot of log10(counts+1) for 19,987 genes expressed in our RNA-seq dataset from dark-grown shoots at 7d post-stratification; the black dot indicates *LTP2*. In Col-0 wild type, *LTP2* was the 7th most highly expressed gene. **C**. Scatterplot of hypocotyl length vs orientation for each individual, measured at 7d post-stratification (MS+1% sucrose); little correlation was found; indicated is Spearman ρ and p-value.

**Figure S2. A**. Boxplots of hypocotyl length, at 7d post-stratification on MS+1% sucrose, for all 5 replicates from Figure 1, showing similar distributions. Replicates 1-3 are from different seed batches, replicates 4-5 are the same seed batch done on different days. **B**. (top) Time-dependent decrease in the Pearson correlation coefficient between hypocotyl and root length at 3, 4, 5 and 7d post-stratification (MS+1% sucrose). Values are similar for *ltp2* and Col-0 supporting similar timing and uniformity of germination in *ltp2*. **C**. Scatterplot of hypocotyl length vs seed area (arbitrary units), showing little correlation; indicated is the Spearman ρ and p-value.

**Figure S3. A**. Hypocotyl (top) and root length (bottom) as a function of time since stratification in Col-0 seedlings grown on MS (open circles) or MS+1%sucrose (filled circles). B. Boxplots comparing the distribution of individual hypocotyl/root ratios for Col-0 seedlings grown on MS (open circles) and MS+1%sucrose (filled circles). C. Boxplots of hypocotyl length for seedlings grown on MS+1% sucrose on either vertical plates (shoot in contact with the media) or horizontal plates (only the roots contact the media); shows that direct sucrose uptake from the shoot is involved in sucrose-induced hypocotyl elongation.

**Figure S4.** Mannitol improves hypocotyl elongation of sucrose-grown *ltp2* seedlings. **A.** Density lines of the distributions of hypocotyl length for Col-0 (grey), *ltp2-1* (blue) and *ltp2-2* (green) seedlings grown on either MS+1% sucrose or MS+1%sucrose+29mM mannitol for 7 days post-stratification.The x- and y-axis range is the same in all graphs. **B**. Dosage-response curves of mean hypocotyl length (y-axis) at 7d post-stratification relative to increasing amounts of mannitol (0-150mM). **C**. Coefficient of variation of hypocotyl length for the distributions shown in **A. D.** Boxplots showing the hypocotyl/root ratio of every seedling shown in **A.**

**Figure S5.** *ltp2* hypocotyls show wild-type-like sucrose-induced responses. **A.** Hypocotyl length as a function of time since stratification in Col-0, *ltp2-1* and *ltp2-2* seedlings grown either on MS (open circles) or on MS+ 1%sucrose (filled circles). **B.** Boxplots comparing the distribution of individual hypocotyl/root ratios for Col-0, *ltp2-1* and *ltp2-2* seedlings grown on MS (positions 1, 3 and 5) and MS+1%sucrose (positions 2, 4 and 6). **C.** Lugol staining of Col-0 (WT), *ltp2-1* and *ltp2-2* etiolated seedlings grown for 7d after stratification on MS (left), MS+1%sucrose (center) or MS+1%glucose. Seedlings were stained for 2min with undiluted lugol solution, rinsed, and left in distilled H_2_O for ∼1hr until imaging. **D.** Boxplots show the distribution of hypocotyl lengths for etiolated Col-0 (WT) and *ltp2-1* seedlings grown for 7d after stratification on either MS+1% sucrose (left, no fill) or MS+2% sucrose (right, filled). Inset (top, right) shows the mean per genotype and condition, and the ratio of the means (2% suc /1% suc) in each genotype.

**Figure S6. A**. Similar numbers of reads (>40 Mio) were obtained for each of the 8 RNA-seq samples. **B**. **left:** Correlation matrix for all 8 x 8 sample comparisons; **right:** Pearson correlation coefficients between replicates are all > 0.98 (see top right corner). **C**. Boxplots of log10(counts) for all 8 samples showing no major difference in count distributions between Col-0 and *ltp2*. **D**. (left) Venn-diagram showing the overlap in DEGs (black; 11) between *ltp2*-L vs *ltp2*-S and Col-L vs Col-S; (right) Fold-change and p-value for the 10 DEGs common to long x short comparisons in Col-0 and *ltp2*. **E**. Boxplot of log10(counts) for all 19,983 genes in our dataset; orange circles: 10 genes from (**D**). **F**. Distance in log10 of the number of DEGs plotted against distance in mean hypocotyl length (7d post-stratification, MS+ 1%sucrose) for each of four sample pairwise-comparisons. **G**. Venn-diagram showing the overlap in DEGs (gray; 474) between the comparison *ltp2*-L vs *ltp2*-S and Col-S vs *ltp2*-S. H. GO term enrichments (fold-enrichment >= 2, p-adj < 0.05) for the set of DEGs unique to Col-S vs *ltp2*-S (1164; see **G**).

**Figure S7.** A. Tree depicting the aminoacid similarity between PR14/Type I LTP proteins. **B**. Plot shows all the Type I LTPs expressed in our RNA-seq dataset of dark-grown shoots (x-axis) and the contribution of each (in %; y-axis) to the subfamily total in these samples. For this analysis, the counts from all four Col-0 samples and all four *ltp2* samples were averaged. **C**. Plot shows the gene expression rank of *LTP2* and *LTP1* relative to all 26,372 genes, across a curated dataset of 56 Arabidopsis RNA-seq samples (see Methods); samples are ordered from highest to lowest; open circle is *LTP1*; filled circle is *LTP2*. In 41 out of 56 samples *LTP1* was higher than *LTP2*, indicating that *LTP1* is the dominant paralog by expression level. **D**. The *LTP1/LTP2* expression ratio is fairly constant across individuals, regardless of their hypocotyl length; shown is the relation between hypocotyl length (at 7d post-stratification on MS+1% sucrose) and the log10(*LTP1/LTP2*) expression ratio for individual shoots. **E**. The *LTP1/LTP2* expression ratio changes during development; dark-grown seedlings have the lowest ratio compared to all other tested samples (light-grown seedlings, roots, leaves and flowers). **Figure S8.** *LTP1* is upregulated across individual *ltp2-1* seedlings. **A.** Scatterplot comparing *LTP1* (y-axis) with *LTP2* expression (x-axis) in individual etiolated shoots grown for 7d post stratification on MS+1%sucrose. **B.** Stacked bar chart depicting the percentage of *LTP1* expression (white) or *LTP2* expression (black) in individual etiolated shoots sorted by increasing hypocotyl length (left to right) and harvested 7d post stratification. **C.** Scatterplot of the relationship between hypocotyl length (y-axis) and *LTP1* expression (x-axis) for a set of wild-type (Col-0, grey) and *ltp2-1* (blue) individual seedlings collected at 7d post stratification (only shoots were used for RNA extraction). Dashed line denotes the line of regression for each genotype. **D.** Scatterplot of the relationship between hypocotyl length (y-axis) and *LTP2* expression (x-axis) for a set of wild-type (Col-0, grey) and *ltp2-1* (blue) individual seedlings collected at 7d post stratification (only shoots were used for RNA extraction). Dashed line denotes the line of regression for each genotype. **E.** Scatterplot of the relationship between hypocotyl length (y-axis) and the log10 of the LTP1/LTP2 expression ratio (x-axis) in wild-type (Col-0, gray, left) and ltp2-1 (blue, right); data are from the same individuals shown in **C** and **D**.

**Figure S9.** *LTP1* and *LTP2* expression is low in the double *ltp1 ltp2* seedlings. Shown are the separate expression levels of *LTP1* (open circles) and *LTP2* (filled circles) used to calculate the *LTP1*/*LTP2* ratio shown in Figure 4D. Each dot represents a bulk sample of about 30 dark-grown shoots collected 7 days post stratification.

**Table S1-S4**. Differential expressed genes between Col-L and Col-S (Table S1), *ltp2*-L and *ltp2*-S (Table S2), Col-L and *ltp2*-L (Table S3), Col-S and *ltp2*-L (Table S4).

**Table S5**. Differentially expressed auxin-responsive genes not shared between Col-S vs *ltp2*-S and *ltp2*-L vs *ltp2*-S.

**Table S6**. Cuticle-related genes from Li-Beisson et al., 2013 expressed in our RNA-seq dataset.

**Table S7**. List of primers used in genotyping and qRT-PCR.

**Table S8**. List of publicly available RNA-seq samples used to establish global expression patterns for *LTP1* and *LTP2*.

