## Supplementary figures and images for "*LTP2* hypomorphs show genotype-by-environment interaction in early seedling traits in *Arabidopsis thaliana*"

### FigS1

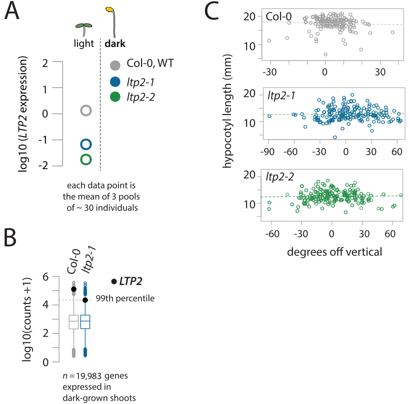

### FigS2

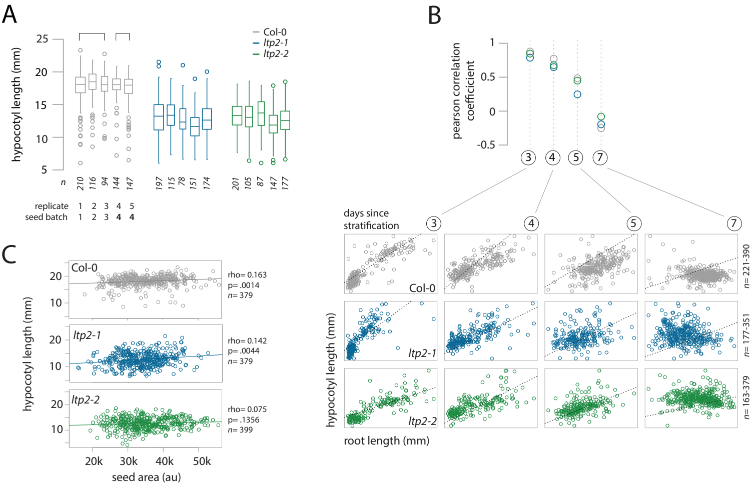

### FigS3

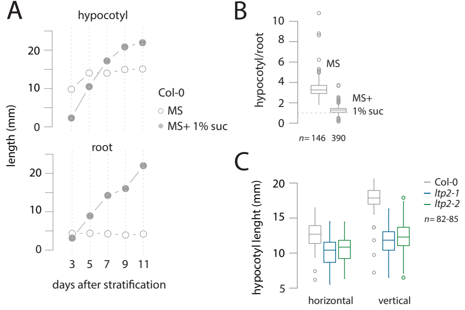

### FigS4

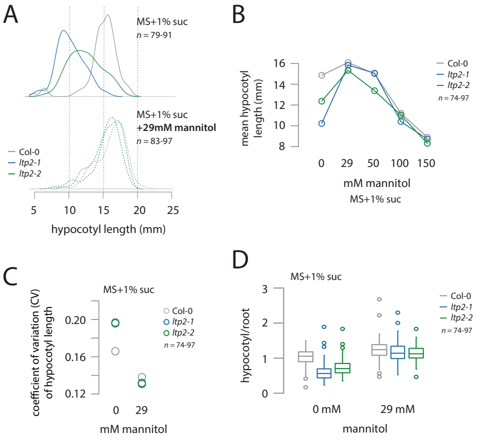

### FigS5

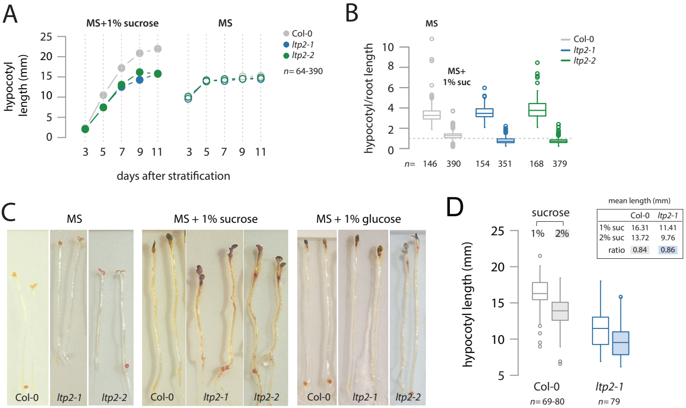

### FigS6

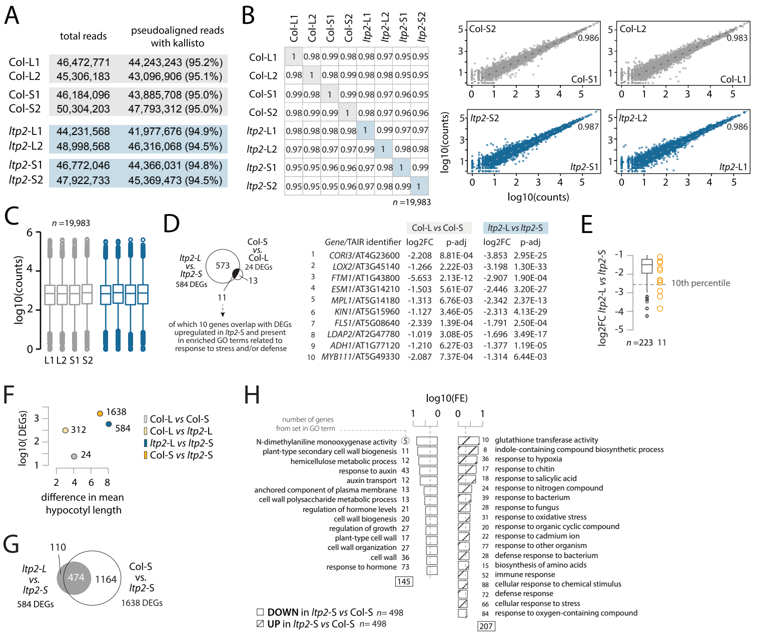

### FigS7

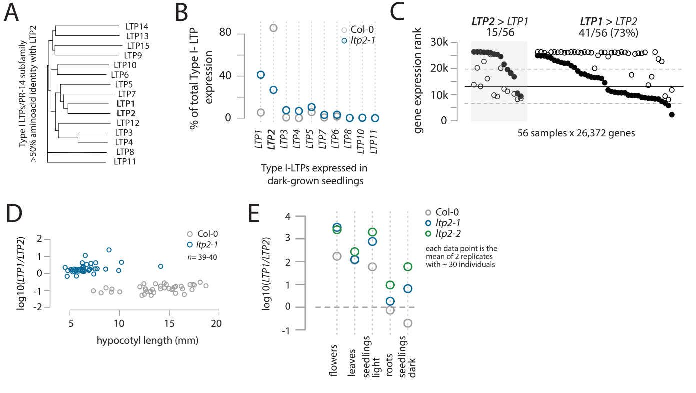

### FigS8

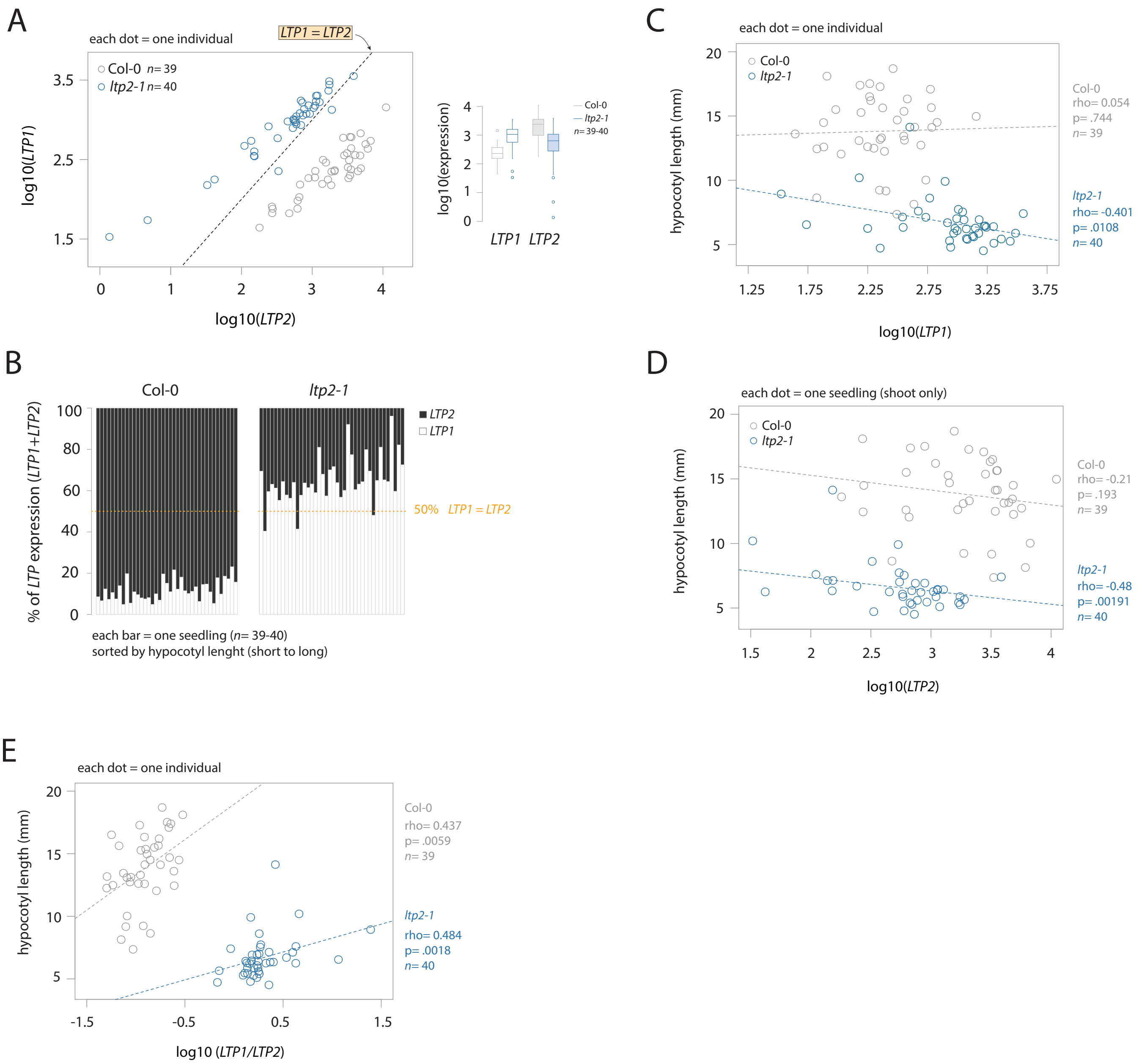

### FigS9

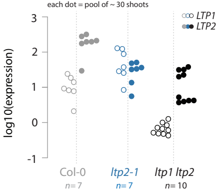
